# Chromosome-level genome assembly and annotation of the threatened marbled teal (*Marmaronetta angustirostris*)

**DOI:** 10.64898/2026.05.12.723956

**Authors:** Joaquín Ortego, Raquel López-Luque, Niclas Backström, Andy J. Green

## Abstract

The marbled teal (*Marmaronetta angustirostris*) is a widely distributed but declining waterfowl species, classified as Near Threatened globally and Critically Endangered in Spain. Despite ongoing conservation actions, including *ex situ* management and population reinforcement programmes, the genomic consequences of long-term captivity, inbreeding, and patterns of functional genetic variation remain unknown due to the absence of a species-specific reference genome. Here, we present the first chromosome-level genome assembly for this species. The genome was generated using PacBio HiFi long reads and Omni-C data, yielding a 1.15⍰Gb assembly with a scaffold N50 of 76.95⍰Mb. A total of 97.16% of the assembly was anchored into 36 chromosome-scale scaffolds, including the Z and W sex chromosomes. BUSCO analysis recovered 99.2% of conserved avian genes. Gene prediction was performed using both ab initio and homology-based strategies, resulting in 16,048 protein-coding genes. This resource provides a foundation for genome⍰wide analyses of inbreeding, demographic history, and adaptive variation, and will support evidence⍰based *in situ* and *ex situ* conservation strategies for this threatened species.

## Background & Summary

The marbled teal (*Marmaronetta angustirostris*) is a small, widely distributed duck inhabiting wetlands across the Mediterranean Basin, North Africa, and south⍰west Asia^1^. Despite its broad geographic range, the species has experienced marked population declines in several regions, particularly in the western Mediterranean, where habitat loss, wetland degradation, and historical hunting pressure have led to severe reductions in population size^2,3^. As a result, the species is currently classified as Near Threatened at the global scale^4^ and Critically Endangered in Spain^3^, which hosts a large fraction of the remaining European population^5,6^.

A recent genetic study characterized the population structure and demographic history of marbled teal, indicating a relatively recent divergence among major biogeographical populations and low levels of genetic differentiation^7^. Despite their shallow evolutionary separation, populations from the western Mediterranean and south⍰west Asia were assigned to distinct genetic clusters and exhibit contrasting demographic and genetic patterns, including elevated inbreeding in declining western populations. Importantly, this study also revealed that captive stocks used in Spain for reinforcement programmes exhibit varying degrees of introgression from long-term European zoo lineages derived from a small founder population maintained in captivity for decades. Combined with evidence that captive-bred individuals often show reduced survival^8^ and altered movement behaviour in the wild^9^, this raises important questions about the evolutionary consequences of captive breeding and its implications for reinforcement programmes^7^.

Despite these advances, genomic resources for *M. angustirostris* remain limited. Available data are restricted to mitochondrial sequences and reduced-representation approaches (ddRAD-seq), which provide only partial coverage of genome-wide variation. These methods are insufficient for detecting genome-wide patterns of inbreeding (e.g., runs of homozygosity), quantifying deleterious mutation load, or identifying signatures of selection associated with long-term captive management (e.g., domestication syndromes). In the absence of a species-specific reference genome, analyses must therefore rely on heterospecific resources from closely related waterfowl, particularly within the genus *Aythya* (e.g., *A. fuligula*^10^, *A. baeri*^11^, and *A. marila*^12^). While these assemblies provide a useful framework within Anatidae, interspecific divergence can affect read mapping, gene annotation transfer, and the resolution of downstream analyses. *Marmaronetta* is also a monotypic genus with no previously available genome assemblies, representing a critical gap in genomic resources for waterfowl. This limitation is particularly relevant in a conservation genomics context, where accurate inference of inbreeding, genomic erosion, and functional variation is essential for management decisions. A high-quality reference genome would therefore enable fine-scale detection of inbreeding patterns, characterization of deleterious variation, assessment of genomic signatures associated with captive breeding, and high-resolution reconstruction of historical demographic trajectories by leveraging the spatial distribution of heterozygosity along chromosomes and recombination-informed coalescent signals that are inaccessible to allele-frequency-based analyses derived from reduced-representation data.

Here, we present the first chromosome-level reference genome assembly for the marbled teal, generated using PacBio HiFi long-read sequencing and Omni-C chromatin conformation capture data from a single female individual. The final assembly spans approximately 1.15 Gb, with a scaffold N50 of 76.95 Mb, and 97.16% of the assembled sequence assigned to 36 chromosome-scale scaffolds, including both Z and W sex chromosomes. Genome completeness is supported by the recovery of 99.2% of conserved single-copy avian genes. Comparative synteny analyses reveal a high degree of genomic collinearity between marbled teal and the closely related tufted duck (*A. fuligula*)^10^, with over 93% of the assemblies aligning in one-to-one relationships. In addition, we provide a complete and annotated mitochondrial genome, a genome-wide characterization of repetitive elements, and a comprehensive set of protein-coding gene annotations. This assembly establishes a reference framework for future analyses of inbreeding, demographic history, and genome variation, and supports the integration of genomic data into conservation management strategies. More broadly, it contributes to the growing catalogue of avian reference genomes and provides a basis for comparative genomic studies within Anatidae.

## Methods

### Sample collection

A blood sample was collected from a captive-bred adult female marbled teal, *Marmaronetta angustirostris*, at Zoobotánico de Jerez, Spain (individual ID: AEP7015209010). Blood was obtained by brachial venipuncture by qualified veterinary staff and collected into EDTA-coated tubes, immediately flash-frozen, and stored at -80 °C until genomic DNA extraction and Omni-C library preparation. All procedures complied with Spanish and European legislation governing animal welfare and the use of animals for scientific purposes (Law 32/2007, Royal Decree 53/2013, and Directive 2010/63/EU). According to institutional and national regulations, no specific ethical approval was required because sampling consisted of a minimally invasive veterinary procedure performed during routine animal handling by authorized personnel at Zoobotánico de Jerez. No animals were sacrificed specifically for this study.

### Genome sequencing

High-molecular-weight genomic DNA was extracted from whole blood by Novogene (Cambridge, UK) using a phenol-chloroform-based protocol optimized to preserve long DNA fragments. PacBio HiFi libraries were prepared and sequenced by Novogene on the PacBio Revio platform. A total of 131.3 Gb of PacBio HiFi data were produced, comprising 6.95 M reads, with a maximum read length of 63,436 bp, a mean read length of 18,891 bp, and a read N50 of 19,850 bp. This corresponded to an estimated genome coverage of ∼114×, based on a final assembled genome size of 1.15 Gb. To enable chromosome-scale scaffolding, an Omni-C library was constructed from the same individual and sequenced on an Illumina NovaSeq X Plus platform by the National Genomics Infrastructure (NGI, SciLifeLab, Stockholm, Sweden). A total of 1,545 M paired-end reads (2 × 150 bp) were generated, providing 463.5 Gb of data. In addition, short-read Illumina sequencing was performed by Novogene on a NovaSeq X Plus platform, generating 116.7 Gb of paired-end reads (2 × 150 bp). These data were used for independent validation of the genome assembly accuracy through read mapping and for mitochondrial genome reconstruction.

### Genome assembly and quality assessment

The genome of *Marmaronetta angustirostris* was assembled using PacBio HiFi long reads combined with Omni-C chromatin conformation data. An initial diploid assembly was generated with Hifiasm (v0.21.0-r686), which integrates PacBio HiFi sequencing data with proximity ligation information to generate phased assembly graphs. We used default parameters except for the inclusion of Omni⍰C reads via the --h1 and --h2 options. Primary contigs derived from the Hi⍰C–integrated assembly graph were retained for downstream analyses. Redundant haplotypic sequences and assembly overlaps were removed using Purge_dups (v1.2.6)^13^ to obtain a haploid representation of the genome. For downstream scaffolding and contact map generation, Omni-C reads were mapped to the purged assembly using BWA-MEM2 (v2.2.1)^14^ with Hi-C–specific flags (-5SP) to retain chimeric and non⍰proper alignments. Omni-C data were processed using Pairtools (v1.1.3)^15^, including parsing, sorting, deduplication, and generation of a filtered pairs file following recommended practices for Hi-C and Omni-C data preprocessing. Scaffolding was performed with YAHS (v1.2.2)^16^, which uses chromatin contact information to order and orient contigs. Hi-C contact maps were generated using Juicer Tools^17^ and PretextMap for visualization and quality control (Fig. 1). The resulting scaffolds were manually curated using Juicebox Assembly Tools^17^, correcting misjoins and refining scaffold structure based on Hi-C interaction patterns. The final chromosome-scale assembly was generated using the juicer post pipeline, which integrates manual curation results to produce the final FASTA and AGP files. Telomeric repeats were identified using the Python script FindTelomeres (https://github.com/JanaSperschneider/FindTelomeres), which detects canonical vertebrate telomere motifs (5⍰-TTAGGG-3⍰ and its complement) at the ends of assembled scaffolds. A total of 97.16% of the assembled sequence was assigned to 36 chromosome-scale scaffolds based on Hi-C data, including the Z and W sex chromosomes (Fig. 2). Chromosome nomenclature was assigned based on synteny with the tufted duck (*Aythya fuligula*)^10^ genome, with chromosome-scale scaffolds named according to their homologous counterparts in this reference (Fig. 2). The final assembly achieved a scaffold N50 of 76.95 Mb, with 253 contigs anchored into chromosome-scale scaffolds, and an overall GC content of 41.68% (Fig. 2). Of the 36 chromosome-scale scaffolds, four contained telomeric sequences at both ends, 15 at one end only, and 17 lacked detectable telomeric sequences (Fig. 2). Assembly completeness was assessed using BUSCO (v5.8.2)^18^ with the *aves_odb10* dataset, yielding 99.2% complete (99.0% single-copy and 0.2% duplicated), 0.1% fragmented, and 0.7% missing genes (Fig. 3).

**Fig. 1.**
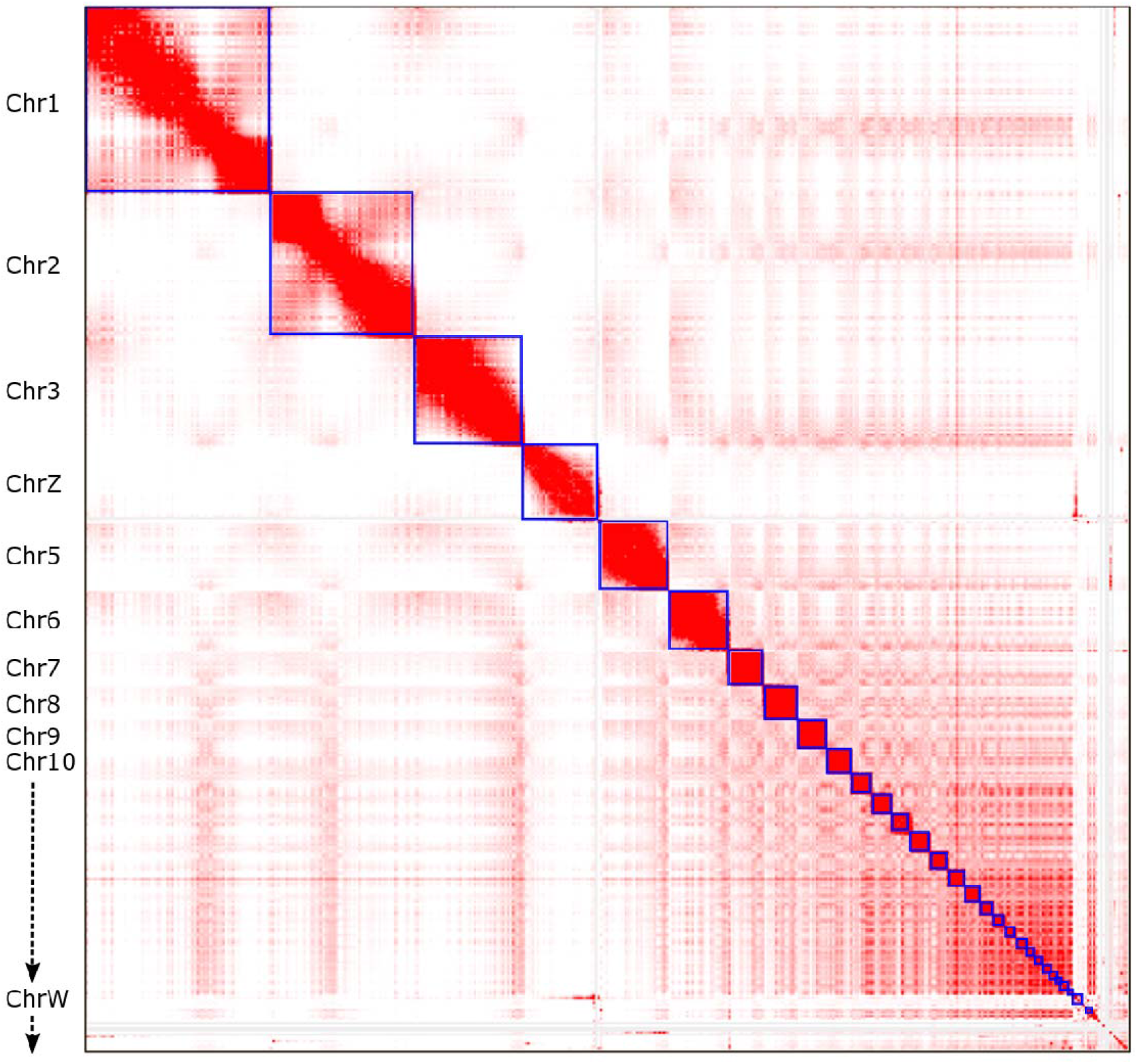
Hi-C contact map of the Marmaronetta angustirostris genome assembly. The contact matrix was generated using Omni-C data. Color intensity reflects interaction frequency between genomic regions, with darker shades indicating higher contact density.

**Fig. 2.**
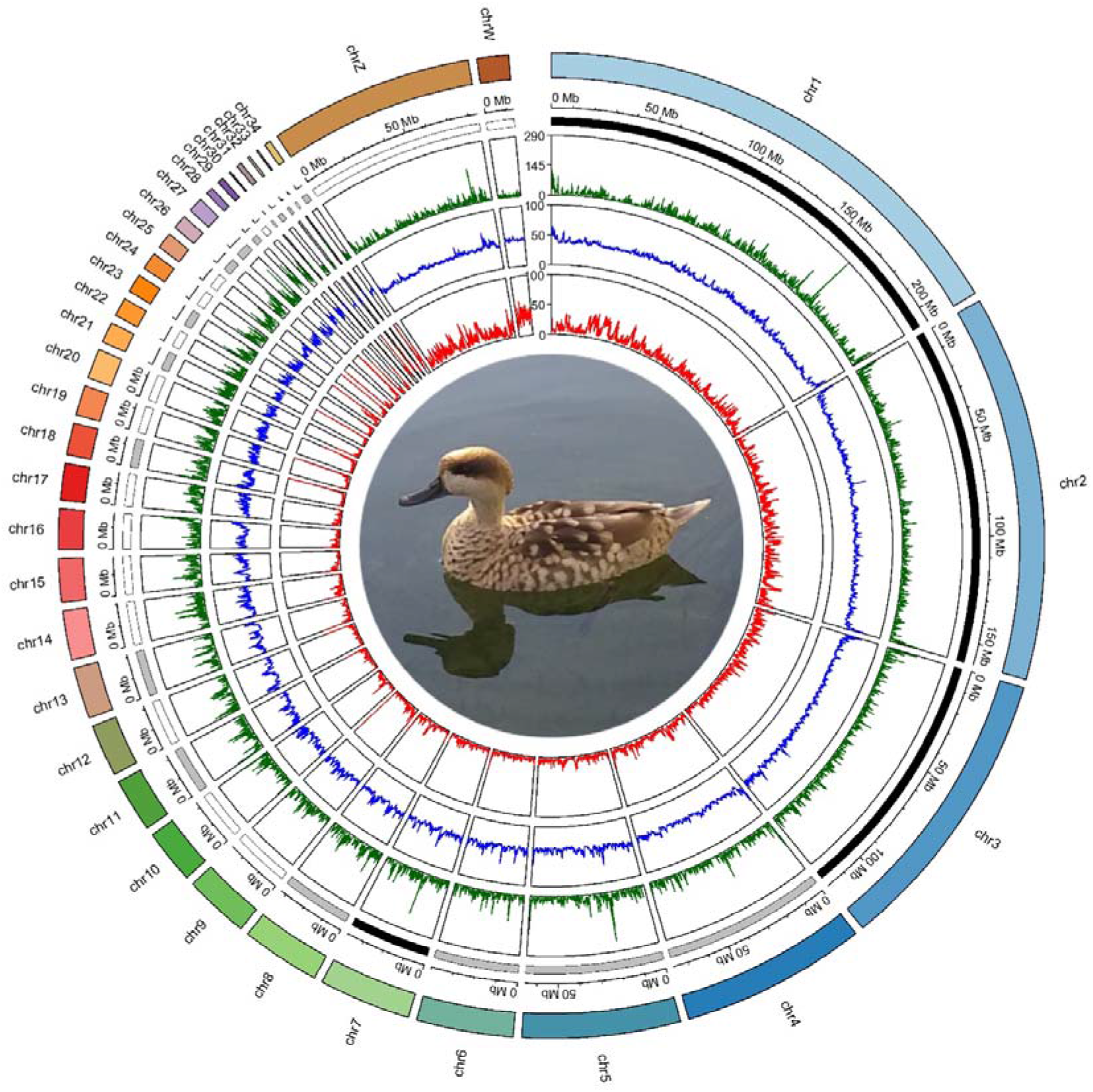
Circos plot of the *Marmaronetta angustirostris* genome. From outer to inner tracks: chromosome ideograms, genomic position scale, telomere presence (black: telomeres at both ends; grey: telomeres at one end only; white: no telomeres), gene density (green), GC content (blue), and repeat density (red), calculated in 100-kb windows. Chromosome nomenclature was assigned based on synteny with the tufted duck (*Aythya fuligula*) genome. The central image shows *M. angustirostris* (photograph by Joaquín Ortego).

**Fig. 3.**
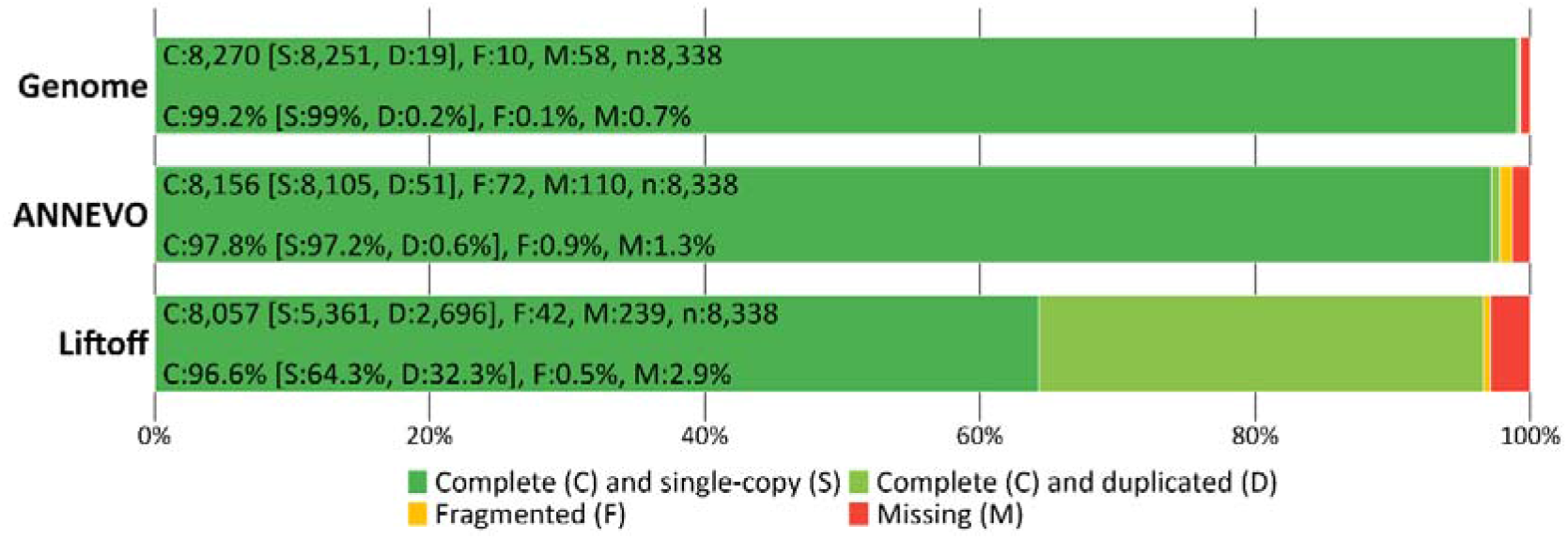
Assessment of Benchmarking Universal Single-Copy Orthologs (BUSCO) completeness in the *Marmaronetta angustirostris* genome assembly and gene annotations. Proportions of complete (single-copy and duplicated), fragmented, and missing BUSCOs are shown for the genome assembly (genome mode) and for gene predictions generated using ANNEVO and Liftoff (protein mode). Analyses were performed against the *aves_odb10* database. Values within each plot indicate the absolute number and percentage of BUSCOs in each category (C: complete, S: single-copy, D: duplicated, F: fragmented, M: missing; n: total number of orthologs searched).

### Mitochondrial genome assembly and annotation

We assembled the mitochondrial genome from Illumina paired-end reads using an iterative mapping approach implemented in MITObim v1.6 (v1.9.1)^19^. Prior to assembly, paired-end reads were interleaved using SeqTK (v1.4, https://github.com/lh3/seqtk) to generate a single read pool. The assembly was initiated using the mitochondrial genome of the tufted duck^10^ as a reference bait, and iterative mapping was performed for 40 cycles using default mismatch parameters. The final consensus sequence was extracted from the last iteration and converted to a single-line FASTA format for downstream processing. To confirm circularity, the consensus sequence was self-aligned using BLASTn (v 2.15.0+)^20^ and MUMmer (v4.0.1)^21^. Terminal overlaps were identified and trimmed to produce a circularized mitochondrial genome. The circularized genome was annotated using the MITOS2^22^ web server under the vertebrate mitochondrial genetic code. Based on the annotation results, the genome was reoriented to start at the tRNA-Phe (trnF) gene, the conventional starting position in avian mitogenomes. The re-oriented sequence was subsequently re-annotated with MITOS2 to confirm gene boundaries and gene-order consistency. The final mitochondrial genome was 16,628 bp in length, which falls within the typical size range reported for avian mitogenomes. The ND3 gene was annotated by MITOS2 as two adjacent fragments with a short overlap, a pattern that has been previously reported in avian mitochondrial genomes and is commonly attributed to lineage⍰specific sequence features of this gene^23,24^. The final mitogenome contains a complete set of 37 mitochondrial genes, including 13 protein-coding genes, 22 tRNAs, and two ribosomal RNA genes, consistent with the canonical organization of vertebrate mitochondrial genomes (Fig. 4).

**Fig. 4.**
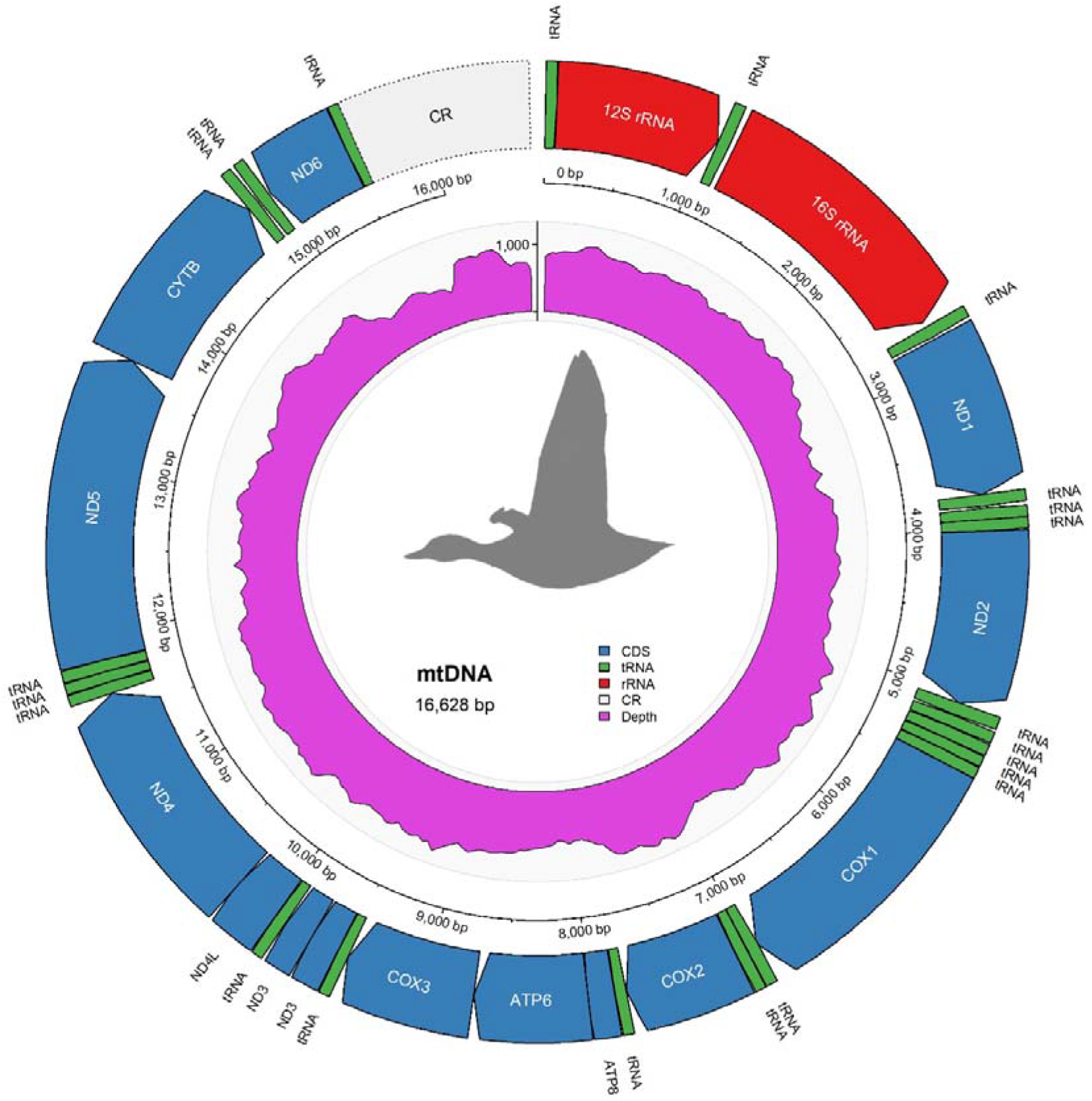
Circular representation of the mitochondrial genome of *Marmaronetta angustirostris*. From outer to inner tracks: annotated genes, genomic position scale, and sequencing depth coverage (shown in pink) calculated in fixed genomic windows of 50 bp. Protein-coding genes (blue), tRNAs (green), rRNAs (red), and the control region (light grey) are shown according to their genomic positions.

### Synteny analysis

Whole-genome synteny between the *Marmaronetta angustirostris* assembly and the tufted duck (*Aythya fuligula*, GCF_009819795.1)^10^ reference genome was assessed using the NUCmer aligner implemented in MUMmer (v4.0.1)^21^ with default parameters. The resulting alignments were processed to calculate the proportion of aligned bases per query scaffold and to assign each scaffold to its corresponding reference chromosome. A total of 93.52% of the *M. angustirostris* genome aligned to the *A. fuligula* reference genome based on one-to-one filtered alignments, with most chromosomes showing >95% sequence alignment. This high proportion of shared sequence reflects strong overall macrosyntenic collinearity between both genomes. Most chromosomes showed clear one-to-one correspondence, supporting the high structural accuracy of the assembly (Fig. 5). Chromosome size comparisons further indicated a generally conserved genome architecture, with only moderate expansions or contractions relative to the reference. Most chromosomes differed by less than ∼0.5 Mb, suggesting limited large-scale structural variation. However, some chromosomes showed more pronounced differences, including expansions in Chr6 and Chr27, and contractions in others (e.g., Chr17, Chr28–30, and the W chromosome), likely reflecting a combination of lineage-specific structural variation and differences in assembly completeness, particularly in repeat-rich regions. Sex chromosomes were identified through synteny patterns. The Z chromosome of *M. angustirostris* showed strong and consistent alignment with the Z chromosome of *A. fuligula* (93.66%), whereas the W chromosome displayed lower alignment (60.93%) and evidence of contraction, consistent with its high repeat content and the known challenges associated with assembling avian W chromosomes. Additionally, some autosomes exhibited partial synteny with the W chromosome, likely reflecting shared repetitive elements. Although most chromosomes exhibited strong synteny, a subset of microchromosomes showed reduced alignment and more fragmented patterns. In particular, Chr31 and Chr34 displayed markedly lower proportions of aligned sequence (44.81% and 11.39%, respectively), and Chr32 also showed reduced alignment (72.80%) compared to the rest of the genome. These chromosomes also showed substantial size discrepancies relative to the reference, especially Chr34, which appears expanded in *M. angustirostris*. Such patterns are consistent with the known difficulties in assembling small, repeat-rich microchromosomes and may reflect both biological variation and residual assembly fragmentation. Overall, the conserved macrosyntenic structure, together with the high proportion of aligned sequence across most chromosomes, supports the completeness and structural reliability of the assembled *M. angustirostris* genome.

**Fig. 5.**
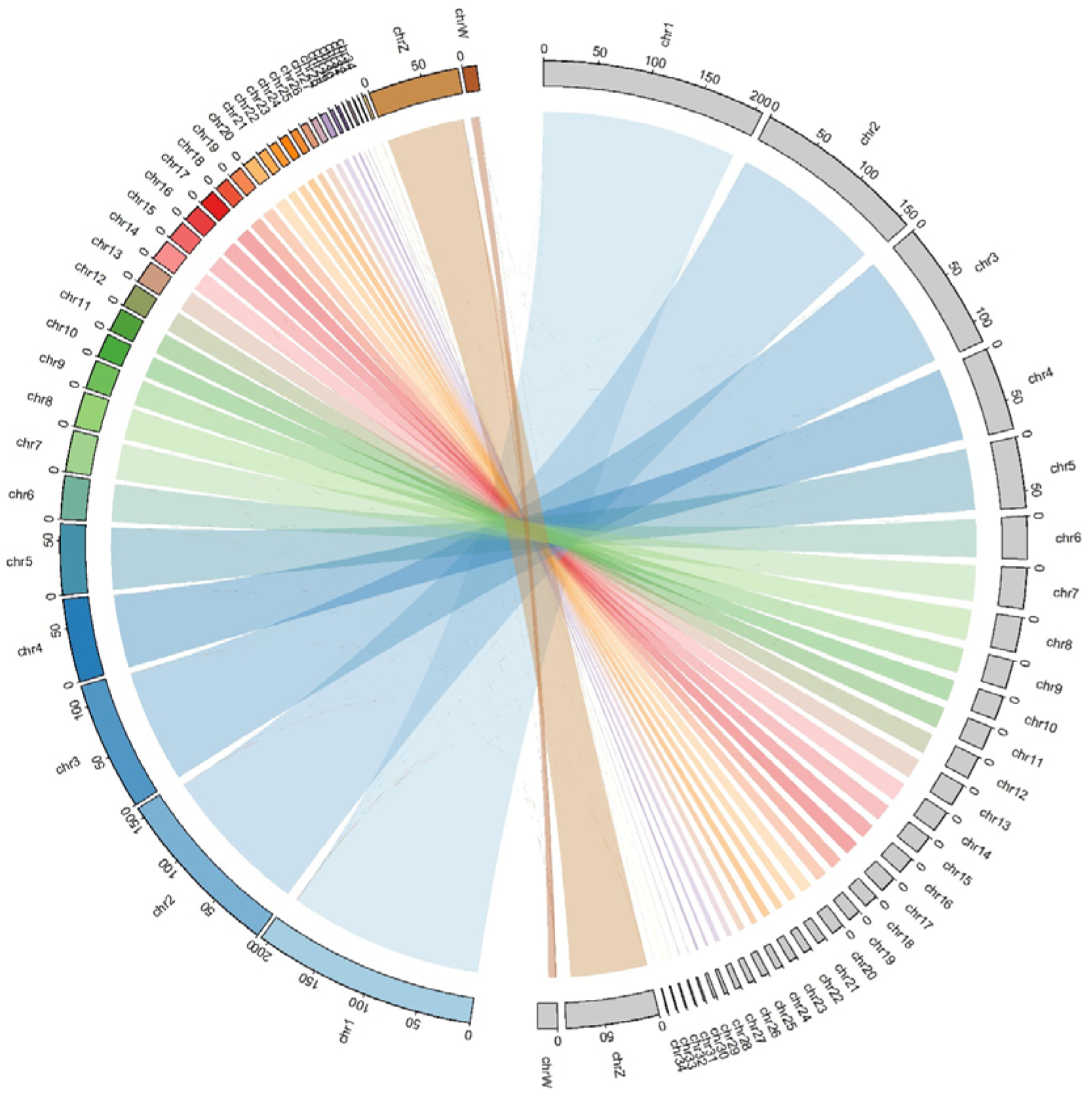
Chromosome-level synteny between *Marmaronetta angustirostris* (colored, left) and *Aythya fuligula* (light grey, right). Links represent one-to-one alignments between homologous genomic regions, illustrating the high degree of macrosyntenic conservation between both genomes. Chromosome sizes are shown in Mb.

### Repeat annotation and masking

Repetitive elements in the genome were identified and masked using a homology-based approach. The final chromosome-level assembly was soft-masked with RepeatMasker (v4.1.5)^25^ using the rmblast search engine (v2.14.1+) and the Dfam database (*CONS-Dfam_3*.*7*)^26^. The species parameter was set to *aves* to improve repeat classification based on curated avian repeat libraries. Soft-masking was performed using the - xsmall option, and repeat annotations were exported in GFF format. Prior to masking, the genome was indexed using BWA-MEM2. Overall, 115.46 Mb of the genome (10.00%) was identified as repetitive sequence (Table 1). Interspersed repeats accounted for 6.47% of the assembly, with retroelements being the dominant class (6.30%). Among these, long interspersed nuclear elements (LINEs) were the most abundant (4.65%), almost entirely represented by CR1-like elements, which are known to dominate avian repeatomes. Long terminal repeat (LTR) elements comprised 1.59% of the genome, while short interspersed nuclear elements (SINEs) were rare (0.06%). DNA transposons constituted a minor fraction (0.13%), and rolling-circle elements were nearly absent (<0.01%). Unclassified interspersed repeats accounted for only 0.03% of the genome, indicating a high level of repeat annotation completeness. In addition to interspersed repeats, simple repeats constituted 3.09% of the genome, while low-complexity regions accounted for 0.41%. Other repeat categories, including small RNAs and satellite DNA, represented minor fractions (<0.1% each). Overall, the repeat landscape is consistent with that of typical avian genomes, which are characterized by relatively low repeat content compared to other vertebrates and a predominance of LINE/CR1 elements. Compared with other Anatidae genomes, the repeat content observed here (10.00%) is slightly lower than that reported for the closely related species *Aythya fuligula* (∼13.0%)^10^ and *Aythya baeri* (∼13.7%)^11^. As in other ducks, LINE elements represent the major component of the repeatome, while SINEs and DNA transposons contribute minimally to genome composition.

**Table 1.** Classification of repetitive elements in the *Marmaronetta angustirostris* genome identified using RepeatMasker.

| Feature | Count | Length (bp) | Percentage (%) |
| --- | --- | --- | --- |
| Genome size | – | 1,154,600,265 | – |
| Bases Masked | – | 115,460,132 | 10.00 |
| SINE | 5,787 | 700,004 | 0.06 |
| Retroelements | 156,454 | 72,767,345 | 6.30 |
| LINEs | 122,066 | 53,714,178 | 4.65 |
| LTR Elements | 28,601 | 18,353,163 | 1.59 |
| DNA Transposons | 9,507 | 1,505,520 | 0.13 |
| Rolling Circles | 70 | 11,178 | 0.00 |
| Unclassified | 2,333 | 396,773 | 0.03 |
| Total Interspersed Repeats | – | 74,669,638 | 6.47 |
| Small RNA | 2,554 | 519,784 | 0.05 |
| Satellites | 570 | 70,353 | 0.01 |
| Simple Repeats | 543,587 | 35,723,007 | 3.09 |
| Low Complexity | 81,317 | 4,699,653 | 0.41 |

### Genome annotation

Genome annotation was performed using complementary *ab initio* and homology-based approaches, in the absence of species⍰specific transcriptomic data. Protein-coding genes were first predicted using ANNEVO (v2.1)^27^, a deep learning-based framework that infers gene structures from evolutionary patterns. The soft-masked genome assembly was used as input together with a pretrained vertebrate model, and gene prediction was conducted using default parameters. In parallel, a homology-based annotation was generated by transferring gene models from *Aythya fuligula* (GCF_009819795.1)^10^ to the assembled genome using Liftoff (v1.6.3)^28^. Alignments were performed requiring a minimum sequence identity of 0.90 and a minimum coverage of 0.95. Both annotation sets were processed using AGAT (v1.4.1)^29^ to standardize GFF3 formatting, remove gene models lacking coding sequences, and filter out genes with coding sequences shorter than 300 bp. After filtering, the ANNEVO annotation comprised 16,048 genes, whereas the Liftoff annotation included 15,244 genes (Table 2). Annotation completeness was assessed using BUSCO (v5.8.2)^18^ in protein mode against the *aves_odb10* lineage dataset. The ANNEVO annotation achieved a completeness score of 97.0% (96.4% single-copy, 0.6% duplicated), with 0.7% fragmented and 2.3% missing BUSCOs (Fig. 3). In comparison, the Liftoff annotation showed a similar level of completeness (96.6%) but a substantially higher proportion of duplicated genes (32.3%), consistent with redundancy introduced during homology-based projection (Fig. 3). Gene structure statistics were broadly consistent between methods, including comparable coding sequence lengths and exon counts per transcript. However, Liftoff annotations exhibited longer average transcript lengths, likely reflecting the transfer of extended untranslated regions (UTRs) from the reference genome and, in some cases, imperfect boundary resolution during projection. Overall, the ANNEVO gene set provided the most reliable representation of the gene space, combining high completeness with low redundancy and internally consistent gene models, and was therefore retained as the primary annotation. The Liftoff results nonetheless supported the overall completeness of the assembly and provided an independent line of evidence for gene content, but were not used as the final gene set due to elevated duplication levels.

**Table 2.** Summary statistics of gene annotation features in *Marmaronetta angustirostris* obtained using ANNEVO and Liftoff.

|  | ANNEVO | Liftoff |
| --- | --- | --- |
| Gene number | 16,048 | 15,244 |
| Average transcript length | 26,387 | 33,869 |
| Average CDS length (bp) | 1,807 | 1,827 |
| Average exons number | 11 | 11 |
| Average exon length (bp) | 159 | 244 |
| Average intron length (bp) | 2,371 | 3,019 |

### Functional annotation

Protein-coding genes were functionally annotated using complementary homology- and domain-based approaches. Predicted protein sequences from the primary annotation (ANNEVO) were aligned against the UniProtKB/Swiss-Prot^30^, UniProtKB/TrEMBL^31^, and NCBI NR databases using DIAMOND (v2.1.8)^32^ in BLASTp mode (e-value < 1e-5), retaining the best hit per query. Domain-based annotation was performed with InterProScan (v5.77-108.0)^33^, enabling the identification of conserved protein domains, families, and functional sites, as well as the assignment of Gene Ontology terms and pathway annotations. Default parameters were used, and Gene Ontology terms were retrieved using the -goterms option. Homology searches identified matches for 95.97%, 99.53%, and 99.59% of proteins in Swiss-Prot, TrEMBL, and NR, respectively. Domain-based annotation detected conserved domains in 99.26% of proteins, with 94.42% associated with Pfam entries and 89.84% assigned at least one GO term. Overall, 16,001 genes (99.71%) were functionally annotated by at least one approach (Table 3), with overlaps among main annotation sources shown in Fig. 6.

**Table 3.** Summary of functional annotation of predicted protein-coding genes in *Marmaronetta angustirostris*.

| Database | Number | Percent (%) |
| --- | --- | --- |
| SwissProt | 15,401 | 95.97 |
| TrEMBL | 15,972 | 99.53 |
| NR | 15,983 | 99.59 |
| Pfam | 15,153 | 94.42 |
| GO | 14,418 | 89.84 |
| InterPro | 15,930 | 99.26 |
| Total | 16,001 | 99.71 |

**Fig. 6.**
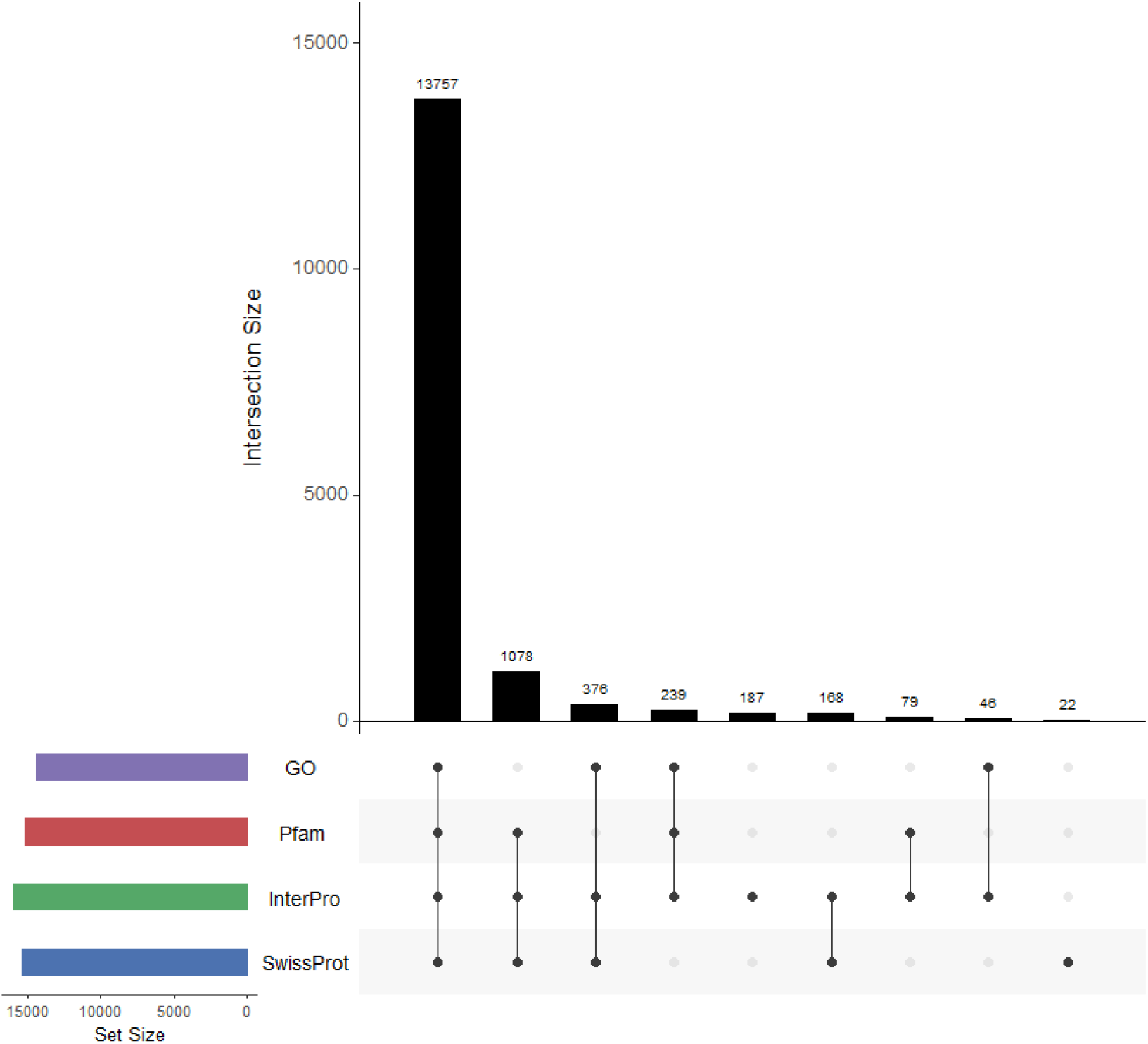
UpSet plot showing the overlap among homology-based matches to UniProtKB/Swiss-Prot and domain-based annotations obtained with InterProScan, including Pfam domains and Gene Ontology (GO) terms.

### Data Records

All raw sequencing data generated in this study were deposited in the NCBI Sequence Read Archive under a single BioProject PRJNA1455455 and BioSample SAMN57366166. PacBio HiFi long reads, Omni-C chromatin conformation capture data, and Illumina short reads are available under accession numbers SRR38198287, SRR38198286, and SRR38198285, respectively. The final genome assembly has been deposited in GenBank under the accession number JBXOSA000000000. Genome annotation files, including structural and functional annotations, as well as repeat annotation results, have been deposited in the Figshare (https://doi.org/10.6084/m9.figshare.32063670).

### Technical Validation

The quality of the input DNA was assessed prior to sequencing using Qubit fluorometry, NanoDrop spectrophotometry, and agarose gel electrophoresis. These analyses confirmed high molecular weight DNA (main gDNA band ≥30⍰kb), appropriate purity, and sufficient concentration for long-read sequencing. Sequencing data quality was evaluated for all platforms. PacBio HiFi sequencing generated highly accurate long reads (read N50: 19.85⍰kb; mean read: 18.89 kb; max length: 63.44 kb), consistent with high-fidelity performance. Illumina short-read and Omni-C datasets exhibited high base quality scores (Q20⍰>⍰98%, Q30⍰>⍰94%), supporting their suitability for genome assembly, scaffolding, and validation. The quality and completeness of the genome assembly were assessed using multiple complementary approaches. Assembly contiguity metrics indicated a highly contiguous genome (scaffold N50: 76.95⍰Mb), with 97.16% of the total assembly assigned to 36 chromosome-scale scaffolds, including both sex chromosomes (Fig. 2). Gene-space completeness was evaluated using BUSCO^18^, recovering 99.2% of conserved avian genes, indicating near-complete representation of the expected gene content (Fig. 3). To further assess assembly accuracy, raw sequencing reads were mapped back to the final assembly. A total of 99.94% of PacBio HiFi reads and 99.46% of Illumina short reads aligned successfully, supporting both base-level accuracy and completeness of the assembly. In addition, potential contamination was evaluated using the NCBI FCS-GX (v0.5.5)^34^ pipeline with the latest genome cross-species database (gxdb; --tax-id 75875), and no contaminant sequences were detected. Chromosome-scale structural accuracy was validated using Omni-C contact maps, which showed strong interaction signals along the main diagonal and well-defined chromosomal boundaries, consistent with correct ordering and orientation of scaffolds (Fig. 1). This was further supported by whole-genome synteny analysis against the closely related species *Aythya fuligula*, with 93.52% of the genome aligning in one-to-one relationships and most chromosomes exhibiting high collinearity, indicating conserved genome structure and supporting the reliability of large-scale assembly (Fig. 5). The mitochondrial genome assembly was independently validated by mapping Illumina paired-end reads, resulting in a high and uniform sequencing depth (mean coverage: 906×; coefficient of variation = 0.15; Fig. 4). This uniform coverage, together with the absence of coverage gaps, supports the completeness and accuracy of the mitochondrial genome assembly. The gene content and organization were consistent with the canonical vertebrate mitochondrial genome (Fig. 4). Finally, genome annotation quality was assessed using BUSCO in protein mode. The *ab initio* gene set achieved 97.0% completeness with low duplication, indicating a reliable and non-redundant representation of the gene space (Fig. 3).

## Code availability

All data processing and analyses were performed using standard bioinformatic tools, following the manuals and recommended protocols of the corresponding software. No custom scripts were developed specifically for the analyses reported here. Software versions and key parameters are reported in the main text, and non⍰default parameters are explicitly indicated where applicable.

## Acknowledgements

We thank the Zoobotánico de Jerez for providing the blood sample from a captive specimen, and íñigo Sánchez García for assistance with sample collection. We are also grateful to Mattias Ormestad (National Genomics Infrastructure, NGI, Sweden) for support during Omni-C library preparation and sequencing, and to Remi-André Olsen (NGI Sweden), Aleix Palahí (Uppsala University), and Erik Gudmunds (Uppsala University) for advice on genome assembly and annotation. We acknowledge the Doñana Singular Scientific-Technical Infrastructure (ICTS-RBD) for access to computational resources, as well as the support provided by the Laboratorio de Ecología Molecular (LEM-EBD) at the Estación Biológica de Doñana (CSIC). The authors acknowledge support from the National Genomics Infrastructure in Genomics Application Stockholm, funded by Science for Life Laboratory, the Knut and Alice Wallenberg Foundation, and the Swedish Research Council, as well as from NAISS/Uppsala Multidisciplinary Center for Advanced Computational Science for assistance with massively parallel sequencing and access to the UPPMAX computational infrastructure. This research was funded by an EU LIFE Project through Fundación Biodiversidad of the Spanish Ministry for the Ecological Transition and the Demographic Challenge (project LIFE19/NAT/ES/000906), by SEO/BirdLife, and by the Severo Ochoa programme (project CEX2024-001498-S, funded by MICIU/AEI/10.13039/50110). N.B. was supported by a project grant from the Swedish Research Council (VR 2019-04791). J.O. was supported by a “Salvador de Madariaga” mobility fellowship (PRX23/00364) funded by the Spanish Ministry of Science, Innovation and Universities, within the State Programme for Talent Promotion and Employability, for a research stay at Uppsala University (Sweden).

## Author contributions

J.O. and A.J.G. conceived the study. R.L.-L. collected the sample and performed the laboratory work. J.O. conducted the bioinformatic analyses, with input from N.B. J.O. wrote the manuscript. All authors reviewed and approved the final version of the manuscript.

## Additional information

**Correspondence** and requests for materials should be addressed to J.O.

## References

1 Salvador, A., Amat, J. A. & Green, A. J. in Birds of the World (eds S. M. Billerman & B. K. Keeney) (Cornell Lab of Ornithology, 2023).

2 Green, A. J. The status and conservation of the marbled teal Marmaronetta angustirostris. IWRB Special Publication 23, 107 (1993).

3 Giménez, M., Botella, F. & Pérez-García, J. M. in Libro rojo de las aves de España (ed N. López-Jiménez) 178–184 (SEO-Birdlife, 2021).

4 BirdLife International. Species factsheet: Marmaronetta angustirostris, <http://www.birdlife.org> (2022).

5 Green, A. J., El Hamzaoui, M., El Agbani, M. A. & Franchimont, J. The conservation status of Moroccan wetlands with particular reference to waterbirds and to changes since 1978. Biol. Conserv. 104, 71–82, 10.1016/s0006-3207(01)00155-0 (2002).

6 Perennou, C., Beltrame, C., Guelmami, A., Tomas Vives, P. & Caessteker, P. Existing areas and past changes of wetland extent in the Mediterranean region: An overview. Ecologia Mediterranea 38, 53–66 (2012).

7 Ortego, J. et al. Demographic and conservation genomic assessment of the threatened marbled teal (Marmaronetta angustirostris). Evol. Appl. 17, 10.1111/eva.13639 (2024).

8 Pérez-García, J. M., Sebastián-González, E., Rodríguez-Caro, R., Sanz-Aguilar, A. & Botella, F. Blind shots: non-natural mortality counteracts conservation efforts of a threatened waterbird. Animal Conservation 27, 293–307, 10.1111/acv.12906 (2024).

9 Pacheco-Guardiola, I. et al. Wetland location and captive breeding influence trans-Mediterranean movements in the endangered marbled duck (Marmaronetta angustirostris). Ibis 168, 245–258, 10.1111/ibi.13442 (2026).

10 Mueller, R. C. et al. A high-quality genome and comparison of short-versus long -read transcriptome of the palaearctic duck Aythya fuligula (tufted duck). Gigascience 10, giab081, 10.1093/gigascience/giab081 (2021).

11 Zhang, L. et al. Chromosome-level genome assembly of the critically endangered Baer’s pochard (Aythya baeri). Sci. Data 10, 176, 10.1038/s41597-023-02063-9 (2023).

12 Zhou, S. Y. et al. A high-quality chromosomal-level genome assembly of greater scaup (Aythya marila). Sci. Data 10, 254, 10.1038/s41597-023-02142-x (2023).

13 Guan, D. F. et al. Identifying and removing haplotypic duplication in primary genome assemblies. Bioinformatics 36, 2896–2898, 10.1093/bioinformatics/btaa025 (2020).

14 Vasimuddin, M., Misra, S., Li, H., Aluru, S. & Ieee. in 33rd IEEE International Parallel and Distributed Processing Symposium (IPDPS). 314–324 (2019).

15 Abdennur, N. et al. Pairtools: From sequencing data to chromosome contacts. PLoS Comput. Biol. 20, e1012164, 10.1371/journal.pcbi.1012164 (2024).

16 Zhou, C. X., McCarthy, S. A. & Durbin, R. YaHS: yet another Hi-C scaffolding tool. Bioinformatics 39, btac808, 10.1093/bioinformatics/btac808 (2023).

17 Durand, N. C. et al. Juicer provides a one-click system for analyzing loop-resolution Hi-C experiments. Cell Systems 3, 95–98, 10.1016/j.cels.2016.07.002 (2016).

18 Simao, F. A., Waterhouse, R. M., Ioannidis, P., Kriventseva, E. V. & Zdobnov, E. M. BUSCO: assessing genome assembly and annotation completeness with single-copy orthologs. Bioinformatics 31, 3210–3212, 10.1093/bioinformatics/btv351 (2015).

19 Hahn, C., Bachmann, L. & Chevreux, B. Reconstructing mitochondrial genomes directly from genomic next-generation sequencing reads-a baiting and iterative mapping approach. Nucleic Acids Research 41, e129, 10.1093/nar/gkt371 (2013).

20 Gertz, E. M., Yu, Y. K., Agarwala, R., Schäffer, A. A. & Altschul, S. F. Composition-based statistics and translated nucleotide searches:: Improving the TBLASTN module of BLAST. BMC Biology 4, 41, 10.1186/1741-7007-4-41 (2006).

21 Kurtz, S. et al. Versatile and open software for comparing large genomes. Genome Biol. 5, R12, 10.1186/gb-2004-5-2-r12 (2004).

22 Donath, A. et al. Improved annotation of protein-coding genes boundaries in metazoan mitochondrial genomes. Nucleic Acids Research 47, 10543–10552, 10.1093/nar/gkz833 (2019).

23 Mindell, D. P., Sorenson, M. D. & Dimcheff, D. E. An extra nucleotide is not translated in mitochondrial ND3 of some birds and turtles. Molecular Biology and Evolution 15, 1568–1571, 10.1093/oxfordjournals.molbev.a025884 (1998).

24 Andreu-Sánchez, S., Chen, W. J., Stiller, J. & Zhang, G. J. Multiple origins of a frameshift insertion in a mitochondrial gene in birds and turtles. Gigascience 10, giaa161, 10.1093/gigascience/giaa161 (2021).

25 Tarailo-Graovac, M. & Chen, N. Using RepeatMasker to identify repetitive elements in genomic sequences. Current Protocols 25, 1–14, 10.1002/0471250953.bi0410s05 (2004).

26 Storer, J., Hubley, R., Rosen, J., Wheeler, T. J. & Smit, A. F. The Dfam community resource of transposable element families, sequence models, and genome annotations. Mobile DNA 12, 2, 10.1186/s13100-020-00230-y (2021).

27 Zhang, P. Y. et al. Highly accurate ab initio gene annotation with ANNEVO. Nat. Methods 23, 740–748, 10.1038/s41592-026-03036-7 (2026).

28 Shumate, A. & Salzberg, S. L. Liftoff: accurate mapping of gene annotations. Bioinformatics 37, 1639–1643, 10.1093/bioinformatics/btaa1016 (2021).

29 Dainat J. AGAT: Another Gff Analysis Toolkit to handle annotations in any GTF/GFF format. (Version v1.4.1). Zenodo. https://www.doi.org/10.5281/zenodo.3552717

30 Bairoch, A. & Apweiler, R. The SWISS-PROT protein sequence database and its supplement TrEMBL in 2000. Nucleic Acids Research 28, 45–48, 10.1093/nar/28.1.45 (2000).

31 Mitchell, A. et al. The InterPro protein families database: the classification resource after 15 years. Nucleic Acids Research 43, D213–D221, 10.1093/nar/gku1243 (2015).

32 Buchfink, B., Reuter, K. & Drost, H. G. Sensitive protein alignments at tree-of-life scale using DIAMOND. Nat. Methods 18, 366–368, 10.1038/s41592-021-01101-x (2021).

33 Jones, P. et al. InterProScan 5: genome-scale protein function classification. Bioinformatics 30, 1236–1240, 10.1093/bioinformatics/btu031 (2014).

34 Astashyn, A. et al. Rapid and sensitive detection of genome contamination at scale with FCS-GX. Genome Biol. 25, 60, 10.1186/s13059-024-03198-7 (2024).

